# VarSome: The Human Genomic Variant Search Engine

**DOI:** 10.1101/367359

**Authors:** Christos Kopanos, Vasilis Tsiolkas, Alexandros Kouris, Charles E. Chapple, Monica Albarca Aguilera, Richard Meyer, Andreas Massouras

**Author notes:** These authors contributed equally to the work (Christos Kopanos, Vasilis Tsiolkas).

## Abstract

**Summary:** VarSome.com is a search engine, aggregator and impact analysis tool for human genetic variation and a community-driven project aiming at sharing global expertise on human variants.

**Availability:** VarSome is freely available at http://varsome.com.

## 1 Introduction

When researchers or clinicians are investigating a specific variant, they may need to check the variant’s coding effect for different transcripts, its genomic location and neighboring variants, the genes it may affect, its population frequency, the function of the associated protein, relevant phenotypes, related literature, clinical studies and pathogenicity, or any of a wide range of data, all of which are spread out across multiple resources, making them difficult to access. Also, most resources holding variant details will typically only accept queries in a particular format, and only handle known variants, making it difficult to retrieve information on the effect of a novel, unknown variant.

Additionally, experts working in their field in hospitals, clinics and laboratories all over the world have each built up their own knowledge over the years but have no easy way of sharing this expertise. There is no simple, user-editable yet centralized way of marking novel variants encountered during a study as pathogenic or benign. The global community’s knowledge, therefore, remains fragmented.

Here, we present VarSome, a search engine for human genomic variation which enables users to look up variants in their genomic context, collects data from multiple databases in a central location and most importantly, aims to enable the community to freely and easily share knowledge on human variation.

## 2 Methods

VarSome includes information from 30 external databases (Supplementary Table 1). VarSome’s database consists of more than 33 billion data points describing in excess of 500 million variants. To deal with this scale of data, we have developed MolecularDB, an extremely efficient data warehouse particularly adapted to genomics and variant data. Its speed is harnessed by our tool, thalia, also written in C++, which maps a variant to a specific genomic location, identifies equivalent variants, the variant type (frameshift, insertion, deletion etc) and its coding effect (if any). The front end is written in HTML5 and JavaScript (React), with the back end implemented in Python 3 (Django) and C++.

## 3 Results & Discussion

VarSome is a search engine for human genomic variation. Users can search by gene name, transcript symbol, genomic location, variant ID or HGVS nomenclature (2). VarSome can also parse single lines from VCF files to look up the variant they describe. The results are not limited to known variants, any variant of any length may be entered. The Examples page at https://varsome.com/examples gives a full list of the ways VarSome can be queried. Finally, VarSome can easily be embedded into other web-sites, and has already been integrated into Variant Validator (3). If the query is a gene or transcript, the results will show the gene’s the official name, links to external databases, a short description of the gene product’s function, as well as any medical conditions associated with it. For variants, VarSome displays a summary at the top of the results page with the rsID (if any), type, location, HGVS notation and, for indels, a list of all equivalent variants (variants that produce the same genotype). The variant’s genomic context is displayed in a custom-built genome browser. If the variant falls within a gene, the browser will display its exonic structure, multiple transcripts, and regions of interest (such as protein functional domains, binding sites etc.)https://varsome.com/examples gives a full list of the ways VarSome can be queried. Finally, VarSome can easily be embedded into other web-sites, and has already been integrated into Variant Validator (3). If the query is a gene or transcript, the results will show the gene’s the official name, links to external databases, a short description of the gene product’s function, as well as any medical conditions associated with it. For variants, VarSome displays a summary at the top of the results page with the rsID (if any), type, location, HGVS notation and, for indels, a list of all equivalent variants (variants that produce the same genotype). The variant’s genomic context is displayed in a custom-built genome browser. If the variant falls within a gene, the browser will display its exonic structure, multiple transcripts, and regions of interest (such as protein functional domains, binding sites etc.) retrieved from UniProt (1) and nearby structural variants from ClinVar (7). Additionally, the browser displays any other known variants in the same genomic region (Figure 1).

**Figure 1:**
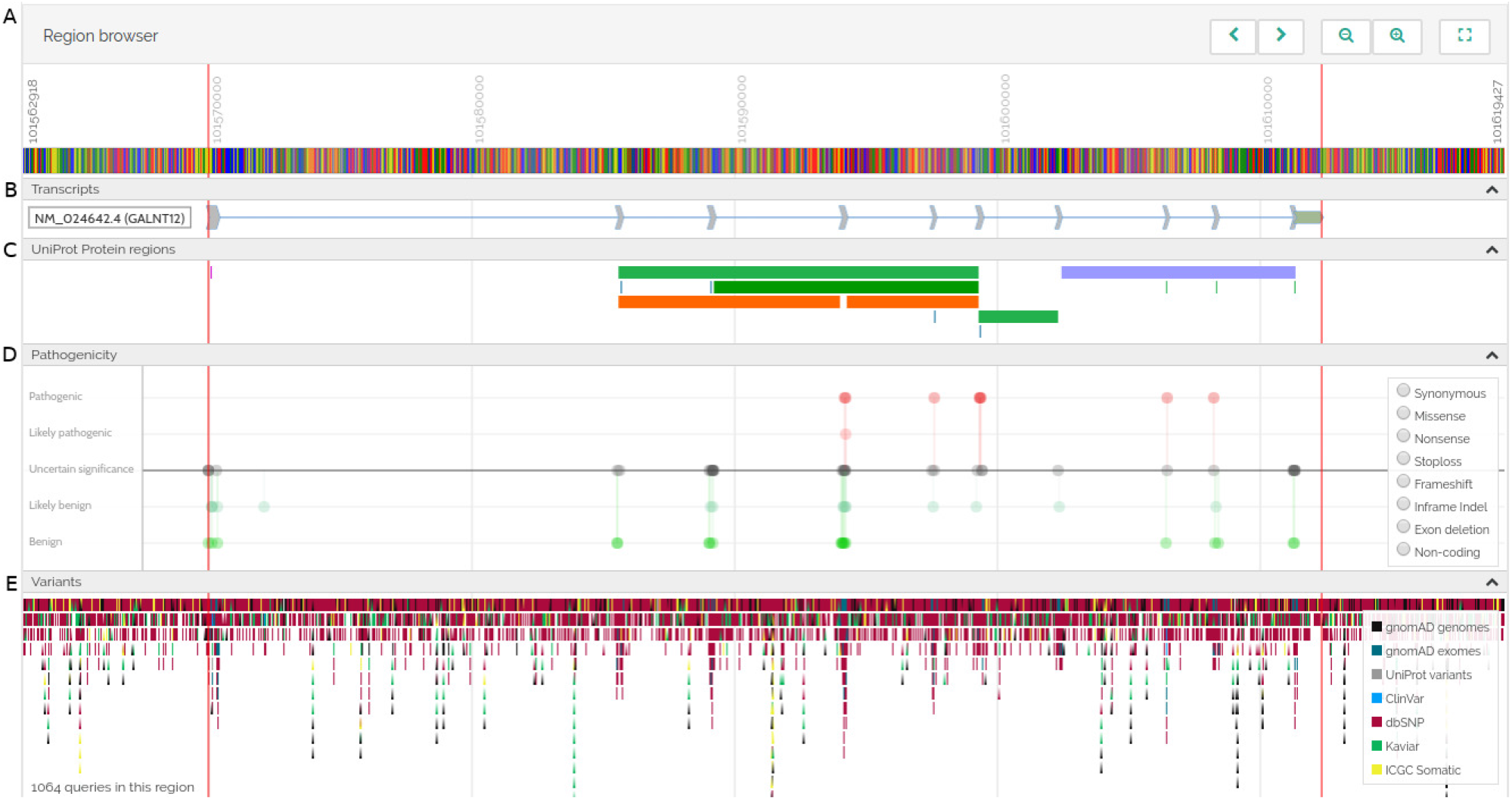
VarSome genome browser. **A)** Sequence (zooming in shows individual base pairs) and position.**B)** Transcripts’ exonic structure and orientation. **C)** Regions of interest in the protein (binding sites, functional domains, etc.) taken from UniProt. **D)** Lollipop graphs indicating the pathogenicity of known variants in the region. **E)** Known variants in the region.

Variant pathogenicity is reported using an automatic variant classifier that evaluates the submitted variant according to the ACMG guidelines (11), classifying it as one of “pathogenic”, “likely pathogenic”, “likely benign”, “benign” or “uncertain significance”. Population frequency data are taken from gnomAD (8), Kaviar3 (4) and ICGC Somatic (5); pathogenicity predictions from dbNSFP (9), which compiles prediction scores from 20 different algorithms, and DANN (10). Clinically relevant information (associated conditions, inheritance mode, publications etc.) are retrieved from the CGD (12), and variants are also linked to any associated phenotypes in the Human Phenotype Ontology (6).

Finally, users can submit their own contributions, linking variants to phenotypes, diseases or articles, and can make their own pathogenicity assessments. These community annotations are linked to the submitting user’s profile and shown to any future users who visit the variant. VarSome also provides short permanent web links to each individual variant, allowing users to easily share their findings or refer to the variant in publications.

## 4 Conclusion

VarSome is both a powerful annotation tool and search engine for human genomic variants, and a platform enabling the sharing of knowledge on specific variants. Since its initial release in May 2016, VarSome has grown to 56,000 users from more than 120 different countries. It has already been integrated into other websites, including the Variant Validator (3), and has been used as an educational resource in university lectures. As the community continues to grow, VarSome is becoming an increasingly important knowledge base for human variation. Most importantly, the ability of users to mark variants as pathogenic or benign allows the combined expertise of the community to be organized and shared to the benefit of everyone in the field.

